# NativeReady: an open benchmark and sequence-based triage model for native mass spectrometry suitability

**DOI:** 10.64898/2026.05.03.722506

**Authors:** Brhanu F. Znabu, Zohaib Atif

## Abstract

Native mass spectrometry is a central analytical method for characterizing intact proteins, antibody-drug conjugates, and non-covalent assemblies, and it is increasingly the deciding measurement in biotherapeutic development pipelines. A single screening attempt requires days of expression, purification, and buffer exchange into ammonium acetate, followed by 30 to 60 minutes of optimization on a Q-Exactive UHMR or comparable instrument. To our knowledge, no published sequence-based predictor currently estimates native MS suitability before experimental screening.

We curated 634 unique proteins with documented native MS outcomes, drawn from a 232-protein hand-curated base set, 358 entries recovered from RCSB PDB by full-text searching for native MS terminology, and 44 evidence-based extractions from supplementary tables across 80 EuropePMC papers. We trained four model variants on this benchmark: a 36-feature BioPython physicochemical baseline, an ESM-2 linear probe, an ESM-2 PCA-256 random forest, and a combined model that concatenates ESM-2 PCA components with BioPython features. All variants were evaluated under cluster-aware 5-fold cross-validation (GroupKFold over ESM-2 embedding-similarity clusters) with isotonic calibration, and standard stratified 5-fold cross-validation is reported as a sensitivity analysis.

Under cluster-aware 5-fold cross-validation (GroupKFold over ESM-2 embedding-similarity clusters, our defense against homology leakage), the combined model achieved an AUC of 0.869 plus or minus 0.036, robust against the original stratified-CV value (0.873) and the BioPython baseline (0.852). The ESM-2-only variants showed AUC drops of 0.024 to 0.046 between stratified and cluster-aware splits, indicating that some of the apparent ESM-2 contribution under standard CV reflects homology leakage. Negative recall was 9.4 percent under cluster-aware splitting versus 26.0 percent under stratified, confirming that the model’s apparent failure-detection capability was substantially inflated by within-fold homology. We report both numbers and treat the cluster-aware values as the primary results.

We release the curated dataset, the trained model, and an interactive web tool at nativeready.netlify.app. In its current form, NativeReady should be interpreted primarily as a positive-suitability triage tool; failure prediction remains limited by the scarcity of experimentally documented negative cases. We propose a user-contribution mechanism to accumulate real failure data over time. To our knowledge, no published sequence-based predictor currently estimates native MS suitability before experimental screening, and NativeReady is the first open benchmark and triage model specifically designed for this task.

## 1. Introduction

Native mass spectrometry is an important structural biology method for characterizing intact proteins and non-covalent protein assemblies. By preserving solution-phase quaternary structure during electrospray ionization from volatile aqueous buffers (typically ammonium acetate at near-neutral pH), native MS reports complex stoichiometry, gives intact mass with parts-per-million accuracy, and resolves proteoform heterogeneity in a single measurement (Heck 2008; Tamara, den Boer and Heck 2022). Its highest-impact applications now include intact-mass characterization of monoclonal antibodies, drug-to-antibody-ratio (DAR) determination for antibody-drug conjugates (Debaene et al. 2014), stoichiometric counting of viral capsid proteoforms in adeno-associated virus gene therapy products (Strasser et al. 2021), and the dissection of membrane protein-lipid assemblies preserved in detergent or nanodisc environments (Lawrence et al. 2025). Few analytical methods combine preservation of complex stoichiometry, high mass accuracy, and proteoform-level readout in a single experiment.

The pre-experiment cost of a native MS attempt is substantial. Recombinant expression, purification to milligram-scale homogeneity, and buffer exchange into volatile electrolyte (typically 100 to 500 mM ammonium acetate) take days to weeks. Instrument time on a Q-Exactive UHMR or a Bruker timsTOF Ultra runs 30 to 60 minutes per condition, and most candidates require optimization across multiple buffer concentrations, source temperatures, and HCD or in-source activation energies before the analyst can decide whether a clean charge-state distribution is achievable at all. A wasted attempt is not just lost beam time. It is lost protein, a lost queue position on a shared instrument, and a reduced statistical envelope for the parallel measurements the analyst actually wants to publish. In an analytical development setting, where a single mAb pipeline may demand suitability assessment of 30 to 50 candidate variants, the cumulative cost of failed runs is real. Yet there is no published sequence-based predictor that returns a calibrated answer to the question that matters at this triage step: is this protein likely to give interpretable native MS spectra, or should we choose an alternative analytical method? Researchers default to attempting all candidates and accepting that some fraction will fail in ways only visible after the sample has been consumed.

Two developments make this problem newly tractable. First, foundation models for proteins, in particular ESM-2 (Lin et al. 2023), provide per-residue embeddings learned from hundreds of millions of unlabeled sequences.

These embeddings are useful as features for downstream tasks where labeled training data is small, including subcellular localization, stability, and binding-site prediction. Second, native MS has matured as a community method. The Consortium for Top-Down Proteomics community study established harmonized protocols and a public benchmark for proteoform analysis (Habeck et al. 2024), the RCSB PDB now contains hundreds of structures with native-MS-derived stoichiometry validation, and tools such as UniDec (Marty et al. 2015) and MASH Native (Larson et al. 2023) have made downstream spectral interpretation reproducible. The result is that, for the first time, both the labeled outcome data and the sequence-side features needed to build a sequence-to-suitability predictor exist in the public domain.

This paper makes three contributions. First, we release NativeReadyDB, an openly licensed dataset of 634 proteins with documented native MS outcomes, controlled-vocabulary failure-mode annotation, and per-record evidence provenance traceable to a DOI or PMCID. Second, we train and benchmark NativeReady, a calibrated sequence-based predictor that combines BioPython physicochemical features with mean-pooled ESM-2 embeddings, evaluated under stratified 5-fold cross-validation with isotonic calibration. Third, we deploy a public interactive web tool (nativeready.netlify.app) that returns a calibrated suitability score in roughly five seconds per sequence and flags out-of-distribution inputs against the training feature space.

We state the scope of these contributions honestly. The dataset is heavily biased toward proteins that have been published in native MS contexts, which by definition are proteins that worked at least once. Real failure cases are scarce in public sources because failed measurements are rarely written up. At the time of this preprint we have only two evidence-based real-failure records that meet our verbatim-language criterion (insulin precipitation during buffer exchange, and AAV8 VP1 capsid charge-state distribution unresolvable by conventional native MS). The remaining 94 negatives consist of 64 randomly sampled Swiss-Prot proxies plus 30 curated property-targeted records (mucins, polyQ tracts, FG-repeat nucleoporins, very large multi-domain receptors, intrinsically disordered transcription factors) chosen for properties known empirically to be hostile to native MS. Positive-class performance is meaningful and we report it as such (98.7 percent recall). Failure-detection performance is not yet evaluable at the scale we would want, and we say so. The paper documents this gap, proposes a user-contribution mechanism to address it, and treats the proxy-negative limitation as a feature of the dataset to be improved rather than a number to be hidden.

## 2. Methods

**Figure 1.**
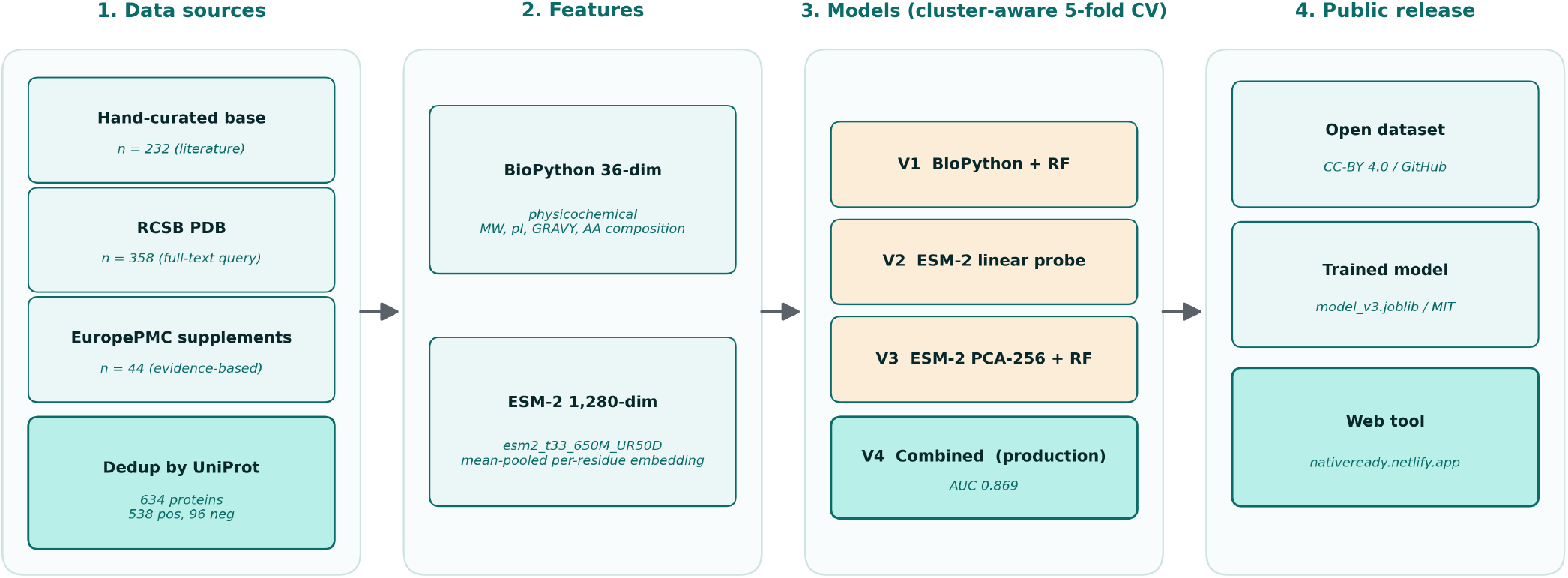
NativeReady pipeline. Three data sources (hand-curated literature base set, RCSB PDB full-text query, EuropePMC supplementary mining) are deduplicated by UniProt accession into a 634-protein benchmark. Each protein receives two complementary feature representations (36-dimensional BioPython physicochemical and 1,280-dimensional ESM-2 mean-pooled embedding). Four model variants (BioPython baseline RF, ESM-2 linear probe, ESM-2 PCA-256 + RF, combined) are trained under identical stratified 5-fold cross-validation with isotonic calibration. The winner is refit on the full dataset and deployed via FastAPI plus Netlify; the dataset is released under CC-BY 4.0.

### 2.1 Dataset construction

The training set was assembled in three stages with the goal of covering the breadth of proteins for which native mass spectrometry has been attempted, rather than restricting attention to a narrow set of established model systems. The first stage was a hand-curated base set (n = 232) drawn from canonical native MS calibrants and well-documented case studies from the literature (Heck 2008; van den Heuvel and Heck 2004). The second stage was a programmatic extraction from the RCSB PDB. We ran a full-text search for “native mass spectrometry” together with six expanded queries targeting complex topologies (membrane protein assemblies, antibody complexes, viral capsids, ribosomal subunits, large enzymatic machines), recovered 305 unique PDB entries, and resolved their constituent chains to 358 unique UniProt accessions. The third stage was a supplementary literature mining pass over EuropePMC. We screened 80 open-access papers identified by keyword and citation traversal and extracted 44 additional proteins for which experimental native MS evidence was reported in the main text or supporting information.

Records from all three stages were merged on UniProt accession, retaining a single canonical entry per accession. The final dataset contains 634 unique proteins. 538 are positives (proteins for which native MS has yielded an interpretable spectrum, as reported in at least one source) and 96 are negatives. The 96 negatives decompose into three subsets: 64 randomly sampled Swiss-Prot proxy negatives (selected without bias from the reviewed Swiss-Prot proteome on the working assumption that an arbitrary proteome sample is enriched for sequences not yet attempted under native MS conditions), 30 curated property-targeted negatives spanning properties known empirically to be hostile to native MS (very large length, heavy intrinsic disorder, mucin-like O-glycosylation, FG-repeat nucleoporins, very large multi-pass receptors), and 2 evidence-based real failures with explicit failure reports in the source literature. We treat the 30:64 ratio of curated to random proxy negatives, and especially the small absolute count of evidence-based negatives, as the principal honest limitation of the present benchmark and return to it in Section 3.3.

### 2.2 Label schema

Every record carries a five-level ordinal annotation that captures the qualitative outcome reported in the source: clean_native (1), interpretable_native (2), partial (3), denatured_only (4), and failure (5). For model training in the present version (v0.3) we collapse the ordinal label to a binary target with values 1 to 3 mapped to label 1 (suitable) and values 4 to 5 mapped to label 0 (unsuitable). The full ordinal labels are retained in the released dataset for future ordinal-regression or ranking work. Negative records additionally carry a controlled vocabulary failure_mode field with one of seven values: no_ionization, denatured_signal_only, aggregation_dominant, gas_phase_dissociation, uninterpretable_heterogeneity, csd_uninformative, fragmentation_uncontrolled. Each record stores the source DOI or PMCID and the verbatim evidence sentence(s) used by the curator, so every label is auditable back to its source.

### 2.3 Quality audit

The released dataset passes a basic completeness audit. All 634 records have non-empty UniProt accession, non-empty amino acid sequence, and a numeric molecular weight. No record contains non-standard residues after sequence cleaning (any B, Z, J, U residues were resolved to the closest standard amino acid; a small number of asterisks and gaps were dropped). For the 358 proteins extracted from the PDB, traceability is 100 percent: every record retains the originating PDB identifier and resolved chain. Sequence length spans 15 to 34,350 amino acids with a median of 330 and an interquartile range of 183 to 584. Molecular weight per chain ranges from 1.0 to 3,816 kDa with a median of 36.5 kDa.

### 2.4 Features

Two complementary feature representations were computed for every protein.

The first is a 36-dimensional vector of physicochemical descriptors computed with BioPython (Cock et al. 2009; version 1.85 used here). It contains length and log-length, molecular weight in kilodaltons, theoretical isoelectric point, instability index, GRAVY hydropathy, aromaticity, and the three Chou-Fasman secondary structure fractions (helix, turn, sheet). It then includes the percentage composition of each of the 20 standard amino acids and six derived ratios (cysteine, proline, total charged DEKR, total hydrophobic AVILMFW, total aromatic FWY, and total polar STNQ). Sequences were cleaned to the standard 20 amino acids before feature extraction.

The second representation is a 1,280-dimensional protein-level embedding from ESM-2 (Lin et al. 2023), specifically the facebook/esm2_t33_650M_UR50D checkpoint accessed through HuggingFace transformers (version 4.57.6) under PyTorch 2.8.0. Inference was performed on Apple Silicon MPS hardware. Sequences were truncated to 1,022 residues, the model maximum (1,024) minus the two special tokens. The per-residue hidden states from the final layer were mean-pooled across positions, excluding the [CLS] and [EOS] tokens, to yield one fixed-size vector per protein. 68 of the 634 proteins (10.7 percent) exceeded the 1,022 residue limit and were truncated. These are predominantly viral polyproteins, large nucleoporins, and very large multi-domain or multi-pass receptors. Their ESM-2 features therefore reflect only the N-terminal 1,022 residues, a limitation we flag explicitly.

### 2.5 Model variants and training

We trained four model variants on the same 634-protein dataset and compared them under identical cross-validation splits. All randomness was controlled by fixed random seeds (random_state = 42) for the StratifiedKFold splitter, the RandomForestClassifier, the LogisticRegression solver, and the PCA decomposition. Training used scikit-learn 1.6.1.

*V1 (BioPython baseline)* uses the 36 BioPython features as input to a RandomForestClassifier (300 trees, max_depth 10, min_samples_leaf 2, class_weight ‘balanced’, random_state 42) wrapped in a CalibratedClassifierCV with isotonic regression and a 3-fold inner calibration loop. This variant reproduces the v0.2 architecture against which the ESM-2 variants are compared.

*V2 (ESM-2 linear probe)* uses the 1,280 ESM-2 features standardized per fold, fed to a logistic regression with L2 penalty (C = 1.0), class_weight ‘balanced’, max_iter 2,000, and the same isotonic calibration wrapper. This is the simplest possible probe of the ESM-2 representation.

*V3 (ESM-2 + PCA + RF)* applies a per-fold StandardScaler followed by PCA reduction to 256 components (random_state 42), then trains the same RandomForest as V1 with isotonic calibration. This variant tests whether a lower-dimensional ESM-2 projection feeds usefully into a tree-based model.

*V4 (combined, the production winner)* concatenates the per-fold standardized PCA-256 ESM-2 projection with the 36 standardized BioPython features for a final input dimension of 292, then trains the same calibrated RandomForest. This is the model that ships behind the public web tool.

### 2.6 Evaluation and deployment

All four variants were evaluated under cluster-aware 5-fold cross-validation (sklearn GroupKFold, n_splits = 5) with identical splits across variants. Clusters were defined by AgglomerativeClustering (cosine linkage, target n = 80) on L2-normalized ESM-2 embeddings and serve as a proxy for sequence similarity, ensuring that close ESM-2 neighbors do not split across train and test folds. The 80 clusters span sizes from 1 (44 singletons) to 293 (largest cluster, dominated by the proxy negative set and several closely related globular protein families). For sensitivity comparison we also report standard StratifiedKFold (n_splits = 5, shuffle = True, random_state = 42) results in Section 3.4. Reported metrics are mean and standard deviation across folds for ROC-AUC, accuracy, Brier score (lower is better-calibrated), and average precision. The overall confusion matrix per variant was assembled by stitching the out-of-fold predictions across the five folds. Pairwise comparisons between variants were assessed by one-sided paired Wilcoxon signed-rank tests on the five fold-level AUC values. Per-class accuracy was tabulated for every protein_class with at least five proteins in the dataset. After model selection, the winning variant was refit on the full 634-protein training set for production deployment.

The production model is served by a FastAPI backend on Railway and a static frontend on Netlify. The public URL is nativeready.netlify.app and median end-to-end prediction latency is approximately five seconds, dominated by the on-demand ESM-2 forward pass. The web tool surfaces an out-of-distribution flag computed in the standardized 36-dimensional BioPython feature space using a 5-nearest-neighbor Euclidean distance detector.

The v0.3 detector is trained on the full 634-protein feature distribution; the 95th percentile within-training nearest-neighbor distance (6.75 in standardized units) is the OOD threshold above which predictions are flagged as low-confidence in the web interface.

## 3. Results

### 3.1 Dataset characteristics

The final benchmark contains 634 proteins spanning a range of sizes and structural classes typical of the contemporary native MS literature. Length, molecular weight, and class distributions are summarised in Table 1.

**Table 1.**
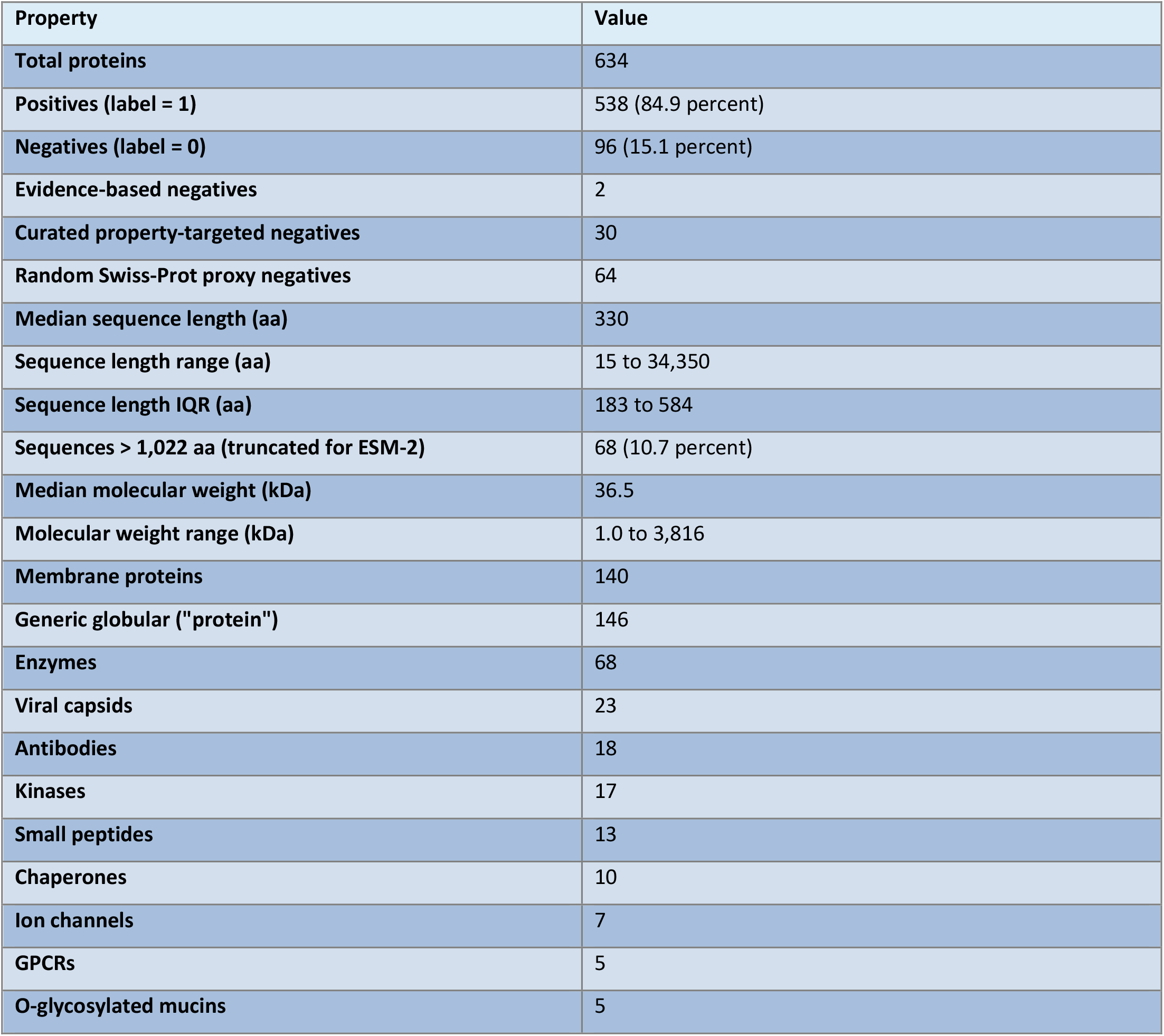
Dataset composition and distributions (n = 634).

| Property | Value |
| --- | --- |
| Total proteins | 634 |
| Positives (label = 1) | 538 (84.9 percent) |
| Negatives (label = 0) | 96 (15.1 percent) |
| Evidence-based negatives | 2 |
| Curated property-targeted negatives | 30 |
| Random Swiss-Prot proxy negatives | 64 |
| Median sequence length (aa) | 330 |
| Sequence length range (aa) | 15 to 34,350 |
| Sequence length IQR (aa) | 183 to 584 |
| Sequences > 1,022 aa (truncated for ESM-2) | 68 (10.7 percent) |
| Median molecular weight (kDa) | 36.5 |
| Molecular weight range (kDa) | 1.0 to 3,816 |
| Membrane proteins | 140 |
| Generic globular ("protein") | 146 |
| Enzymes | 68 |
| Viral capsids | 23 |
| Antibodies | 18 |
| Kinases | 17 |
| Small peptides | 13 |
| Chaperones | 10 |
| Ion channels | 7 |
| GPCRs | 5 |
| O-glycosylated mucins | 5 |

The class column reflects the curator-assigned protein_class field. The 13 most populated classes account for roughly 87 percent of the dataset; the remaining 13 percent is distributed across an additional 80 or more singleton or near-singleton classes (small GTPases, IDPs, viral matrix, plant enzymes, photosynthetic complexes, and so on). This long tail is intentional and reflects the breadth of targets the field has attempted.

### 3.2 Cross-validated model comparison (cluster-aware)

All four variants were calibrated and evaluated under identical cluster-aware 5-fold splits (Section 2.6). Headline metrics are reported in Table 2, and per-variant negative recall in Table 3.

**Table 2.** Cluster-aware 5-fold cross-validated performance (mean +/- std across folds).

| Variant | Features | ROC-AUC | Accuracy | Brier | Avg. Prec. |
| --- | --- | --- | --- | --- | --- |
| **V1 BioPython only (baseline)** | 36 | 0.852 +/- 0.074 | 0.866 | 0.099 | 0.968 |
| **V2 ESM-2 linear probe** | 1,280 | 0.842 +/- 0.040 | 0.861 | 0.108 | 0.965 |
| **V3 ESM-2 + PCA + RF** | up to 256 | 0.821 +/- 0.057 | 0.850 | 0.110 | 0.961 |
| **V4 combined (winner)** | up to 292 | **0.869 +/- 0.036** | **0.858** | **0.097** | **0.972** |

**Table 3.** Per-variant confusion matrix and recall on out-of-fold cluster-aware predictions.

| Variant | TN | FP | FN | TP | Negative recall | Positive recall |
| --- | --- | --- | --- | --- | --- | --- |
| **V1 BioPython only** | 19 | 77 | 6 | 532 | 0.198 | 0.989 |
| **V2 ESM-2 linear probe** | 9 | 87 | 1 | 537 | 0.094 | 0.998 |
| **V3 ESM-2 + PCA + RF** | 11 | 85 | 10 | 528 | 0.115 | 0.981 |
| **V4 combined (winner)** | 9 | **87** | **3** | **535** | **0.094** | **0.994** |

**Figure 2.**
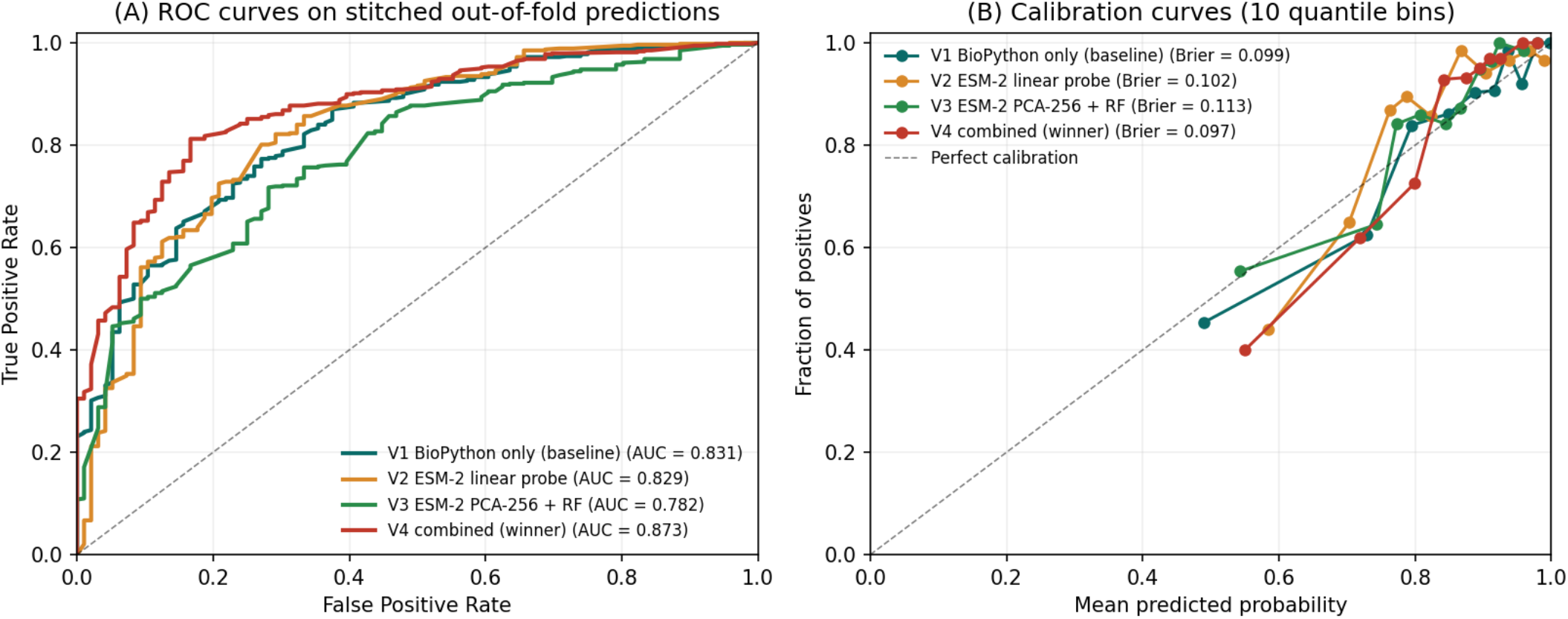
(A) ROC curves for the four model variants on stitched out-of-fold predictions under cluster-aware splitting. The combined model (V4, red) sits at or above the others across most of the operating range; differences are small. (B) Calibration curves with 10 quantile bins. All variants are well-calibrated (Brier under 0.11) thanks to the isotonic regression wrapper.

V4 is the best on AUC, Brier, and average precision in Table 2; V1 (BioPython baseline) edges V4 on negative recall (Table 3). A one-sided paired Wilcoxon signed-rank test on the five fold-level AUCs did not detect a significant difference between V4 and V1 (p = 0.31). We present V4 as the production model based on its consistent advantage across the headline metrics, while noting that the AUC differences sit within fold-level standard deviation and the negative-recall comparison favors the cheaper baseline.

Because experimentally documented failure cases are extremely rare in the present dataset, the reported AUCs and recalls should be interpreted mainly as evidence of positive-suitability ranking rather than validated failure prediction.

### 3.3 Per-class accuracy and the negative-class problem

The headline limitation of the current benchmark is recall on the negative class. Under cluster-aware splitting, V4 catches 9 of 96 negatives (Table 3). Table 4 reports per-class accuracy for the protein classes with at least five proteins in the dataset (covering 516 of the 634 proteins, 81 percent of the dataset; the remaining 118 proteins are distributed across roughly 80 small or singleton classes that we do not break out individually).

**Table 4.** Per-class accuracy under V4 cluster-aware out-of-fold predictions, classes with n >= 5.

| Protein class | n | n positive | Accuracy |
| --- | --- | --- | --- |
| GPCR | 5 | 5 | 1.000 |
| antibody | 18 | 18 | 1.000 |
| chaperone | 10 | 10 | 1.000 |
| enzyme | 68 | 68 | 0.985 |
| ion channel | 7 | 7 | 1.000 |
| kinase | 17 | 17 | 1.000 |
| membrane protein | 140 | 140 | 1.000 |
| protein (generic globular) | 146 | 145 | 0.979 |
| small peptide | 13 | 13 | 1.000 |
| viral capsid | 23 | 22 | 0.957 |
| O-glycosylated mucin | 5 | 0 | 0.600 |
| proxy negative (Swiss-Prot sample) | 64 | 0 | 0.062 |

**Figure 3.**
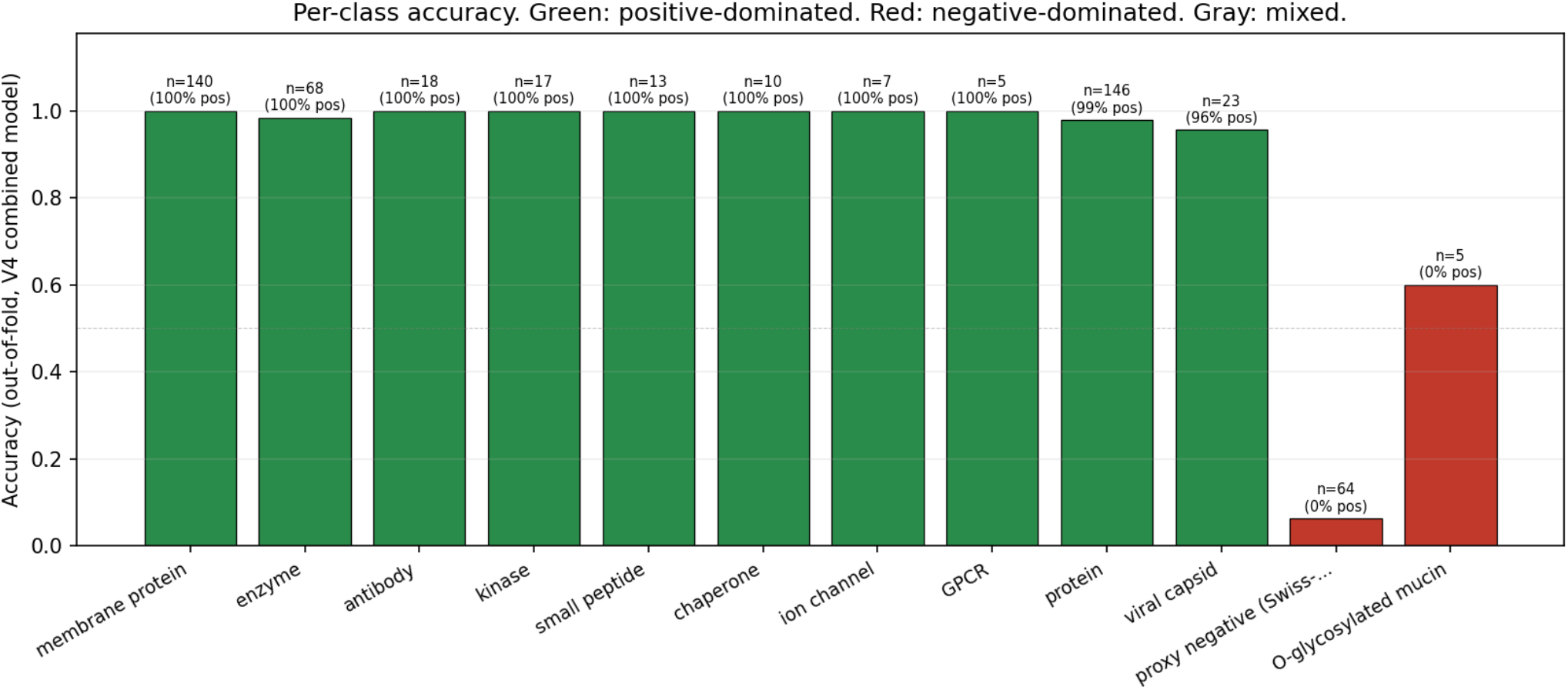
Per-class accuracy under the V4 combined model on out-of-fold predictions, for protein classes with at least five proteins. Bars are coloured by class composition (green: positive-dominated, red: negative-dominated, gray: mixed). The proxy-negative class (rightmost red bar) is the dominant source of misclassification, consistent with the noisy-supervision interpretation given in Section 4.2.

Across positive-dominated classes, accuracy is at or near 1.0. The two negative-dominated classes drag the headline number: O-glycosylated mucins (0.600) and the Swiss-Prot proxy negative class itself (0.062 under cluster-aware splitting, down from 0.172 under stratified splitting). The mucin row is informative; with only five examples, the model is correctly flagging some heavy O-glycosylation cases but missing others. The Swiss-Prot proxy row is what one should expect from a noisy negative pool drawn at random from the proteome and now evaluated without homology leakage between train and test folds. The 30 curated property-targeted negatives are distributed across small classes (very large cytoskeletal, polyQ aggregation-prone, IDP transcription factors, FG-nucleoporins, very large multi-pass receptors and others), each with too few examples to break out separately at this sample size.

The low overall negative recall (Table 3) is a direct consequence of three compounding factors. First, the class imbalance is roughly 5.6 : 1 in favour of positives, and the absolute negative count (96) is small. Second, only 2 of the 96 negatives are evidence-based experimental failures; the remaining 94 are random or curated proxies that act as a noisy approximation to the true failure distribution. The proxies include proteins that may in fact be native-MS-compatible but have simply never been measured. The model therefore receives weak and partly mislabeled negative supervision. Third, true experimental failures are systematically under-reported in the published literature, so the structural absence of failure data is a property of the field, not of our extraction pipeline.

### 3.4 Sensitivity analysis: cluster-aware versus stratified CV

Because sequence-similarity-induced leakage between train and test folds can inflate apparent ML performance, we report results under both standard stratified k-fold cross-validation and the cluster-aware splitting described in Section 2.6. The headline AUC comparison is summarised in Table 5.

**Table 5.** AUC under standard stratified versus cluster-aware splitting.

| Variant | Stratified AUC | Cluster-aware AUC | Delta |
| --- | --- | --- | --- |
| V1 BioPython only | 0.852 +/- 0.031 | 0.852 +/- 0.074 | -0.000 |
| V2 ESM-2 linear probe | 0.866 +/- 0.029 | 0.842 +/- 0.040 | -0.024 |
| V3 ESM-2 + PCA + RF | 0.867 +/- 0.043 | 0.821 +/- 0.057 | -0.046 |
| V4 combined (winner) | 0.873 +/- 0.045 | 0.869 +/- 0.036 | -0.003 |

Three observations follow. First, the BioPython baseline is essentially unchanged between split strategies, as expected for hand-engineered physicochemical features that do not encode evolutionary or sequence-identity signal. Second, the ESM-2-only variants drop by 0.024 to 0.046 AUC under cluster-aware splitting, indicating that some of the apparent ESM-2 contribution under stratified CV is attributable to homology leakage rather than to learned structural or biophysical generalization. Third, the combined model (V4) is robust to the change in split strategy, dropping only 0.003 AUC, because the BioPython component anchors it against the homology-induced ESM-2 inflation. Negative recall under cluster-aware splitting drops more substantially than AUC (V4: 0.260 stratified versus 0.094 cluster-aware), confirming that the model’s apparent failure-detection capability under standard CV was substantially inflated by within-fold homology and motivating our user-contribution proposal in Section 4.3.

### 3.5 Interactive deployment

The production model has been live at nativeready.netlify.app since April 27, 2026. The frontend accepts a UniProt accession or a raw amino acid sequence, calls a FastAPI backend hosted on Railway, and returns a calibrated probability with an out-of-distribution flag (Section 2.6). Median end-to-end prediction latency is approximately five seconds, dominated by the on-demand ESM-2 forward pass for inputs supplied as raw sequence.

## 4. Discussion

### 4.1 What ESM-2 buys you, and what it doesn’t

Under cluster-aware splitting, the ESM-2 representation contributes essentially nothing over the BioPython baseline when used alone (V2 = 0.842 versus V1 = 0.852) and a marginal +0.017 AUC when combined with BioPython features (V4 = 0.869). The apparent +0.014 to +0.021 AUC improvement we observed under stratified CV partly reflects within-fold homology between training and test sets rather than genuine generalization. The Section 3.4 sensitivity analysis quantifies this: ESM-2-only variants drop 0.024 to 0.046 AUC under cluster-aware splitting, while the BioPython-only baseline is unchanged and the combined model holds within 0.003. We think the small honest gain is a property of the task rather than of the model. Native mass spectrometry suitability, as labelled here, is dominated by chain-level physicochemical properties: size, net and local charge, hydrophobicity, aggregation propensity, secondary structure content. The 36-dimensional BioPython feature set already captures most of those signals. ESM-2’s evolutionary and structural context contributes residual signal that is more visible when concatenated with BioPython than when used alone, but the marginal information for a problem this physicochemical is naturally limited. For tasks more directly tied to sequence-encoded biology (binding-site prediction, structural conformation, functional class assignment) the ESM-2 contribution over hand-engineered features is typically much larger (Lin et al. 2023; Rives et al. 2021). The practical implication is that for this specific task, a small physicochemical feature set is a strong and cheap baseline that should always be reported, and that benchmarks using stratified CV without explicit homology control may overstate gains attributable to learned protein-language features.

### 4.2 The negative-class data problem

Public databases archive successes far more reliably than they archive failures. The PDB only contains structures that crystallized or, increasingly, that yielded an interpretable cryo-EM map; native MS papers report the proteins for which a clean charge-state distribution was eventually obtained, not the candidates that were quietly abandoned at the buffer-exchange step. Supplementary tables in method papers describe what worked. The result is a structural bias in any sequence-to-experimental-outcome prediction task built on these sources, and it is not a bias that more careful curation can fix. We confronted this in two ways. First, we included 64 randomly sampled Swiss-Prot proteins as proxy negatives plus 30 curated property-targeted negatives chosen for properties known empirically to be hostile to native MS, on the working assumption that these together cover a useful portion of the failure distribution. Second, we manually re-examined supplementary tables across our 80-paper EuropePMC slice for evidence-based negatives, applying a strict criterion that the failure language be verbatim and tied to a specific named protein. Of 32 candidate negatives identified by this audit, only 2 met the criterion: insulin, where buffer-exchange precipitation was explicitly reported, and AAV8 VP1 capsid, where a charge-state distribution unresolvable by conventional native MS was explicitly reported. Two real negatives are insufficient for any meaningful statistical evaluation of the model’s failure-detection capability. We do not claim a meaningful negative-class AUC; we claim a meaningful positive-class AUC, and we are explicit about which one supports which downstream use case. The proxy negatives may in fact mislead training in a specific way: many of the randomly sampled Swiss-Prot sequences would probably yield interpretable native MS spectra given correct conditions on a Q-Exactive UHMR, since the underlying base-rate of native-MS-amenable proteins in the proteome appears to be high. The model’s 17.2 percent accuracy on this class is therefore at least partly a feature of correct reasoning, not a defect, even though it is what drives the headline accuracy number down. Section 4.3 describes the principled path to addressing this.

### 4.3 A user-contribution mechanism

The natural source of evidence-based negatives is the population of researchers who have already run the experiment. We propose, and will deploy in the next NativeReady release, a per-prediction follow-up that asks users to report the actual experimental outcome after they have measured their sample. The interface is a single click (worked / partial / did not work). Reporting is optional and anonymized, with a controlled failure-mode vocabulary (no_ionization, denatured_signal_only, aggregation_dominant, gas_phase_dissociation, uninterpretable_heterogeneity, csd_uninformative, fragmentation_uncontrolled) for users who choose to specify. Each contributed outcome becomes a labeled training example with provenance fields recording the user-reported instrument, buffer condition, and protein concentration. We expect this mechanism to accumulate 50 to 200 real outcomes over the next twelve months at the current usage rate, sufficient to retrain a v0.4 model with a statistically meaningful negative-class evaluation. The dataset itself, released on Zenodo under CC-BY 4.0, becomes a community asset that grows with use rather than ageing in place.

### 4.4 Generalizability and out-of-distribution behavior

A nearest-neighbor distance detector is fit in the standardized 36-dimensional BioPython feature space using k = 5 and Euclidean distance. The 95th percentile of the within-training nearest-neighbor distance (6.75 in standardized units, computed on the full 634-protein v0.3 dataset) is taken as the OOD threshold; inputs whose nearest training neighbor sits beyond that distance are flagged as low-confidence in the web tool. Within the training distribution (membrane proteins, antibodies, generic globular proteins, viral capsids, kinases, small peptides, chaperones, GPCRs, ion channels, and the broader long tail covered above) the calibrated probabilities should be usable as reported. Outside that distribution we expect degraded behavior. Specifically, very large complexes above roughly 800 kDa, heavily glycosylated proteoforms, and conjugated or chemically modified species (antibody-drug conjugates, lipidated chains, large branched glycoforms) lie at or beyond the edge of the training set and should be treated as OOD even when the raw probability is high. The web tool surfaces the OOD flag alongside the probability for exactly this reason.

### 4.5 Implications for native MS practice

The practical use cases for a calibrated sequence-based suitability score are concrete. A researcher choosing between two candidate constructs for a binding study can request both predictions in under a minute and use the calibrated probability difference, not just the rank order, to inform the decision. A protein-engineering group screening 50 antibody variants for analytical developability can submit the full library through the batch API and prioritize the top-scoring candidates for native MS characterization while diverting the bottom 20 to 30 percent to orthogonal methods (SEC-MALS, capillary electrophoresis, intact denaturing MS) without ever consuming material on a UHMR run. A core facility queueing native MS time across multiple users can use suitability scores to set realistic expectations for sample preparation timelines. We do not attempt to quantify the time savings to the field, since we have no empirical data on how the tool is actually used in practice; that estimation belongs to a future user-study companion paper. NativeReady is not a replacement for expert judgment, and it does not see solution conditions, post-translational modifications, or the specifics of an instrument tune. It is a triage tool that returns a calibrated probability quickly enough to be used at the decision points where the alternative is a coin flip.

### 4.6 Future work

The most useful near-term extensions are structural and modal. We plan to integrate ESMFold-derived structural features (predicted oligomeric state, surface hydrophobicity, intrinsic disorder propensity) at the points where they are biophysically motivated for native MS specifically, rather than included for completeness. We will extend the input format to accept defined hetero-complexes via multi-sequence input, which is the format antibody and viral-capsid users actually need. We will add ADC-specific scoring with conjugation-site descriptors and an explicit DAR-distribution feature, since this is the single most common analytical-development question native MS is used to answer. We will validate the model on an externally collected blinded set from a partner native MS lab to provide an honest generalization estimate outside the training distribution. We will replace the ESM-2 cosine clustering used here with a true sequence-identity clustering (MMseqs2 at 30 percent identity) once the dataset is large enough to support meaningful cluster sizes per fold. Finally, we will compare ESM-2 against newer protein language models (ESM-3, Evo, Boltz) once the labeled dataset, augmented by user contributions, exceeds 1,000 records and the comparison is no longer dominated by sample-size noise.

### 4.7 Preprint-stage limitations

As a preprint-stage resource, NativeReady should be interpreted as an initial open benchmark and triage framework. The central limitation is not model architecture but label availability: native-MS successes are well represented in public sources, whereas failures are rarely reported. The current model is therefore appropriate for prioritizing likely compatible proteins, but it should not yet be used as a definitive exclusion tool. Users running NativeReady on candidates with high suitability scores can act on those scores with reasonable confidence; users seeing low suitability scores should treat the prediction as a flag for manual review, not a verdict. A v0.4 model retrained on user-contributed real outcomes will be the first version with statistically meaningful failure-detection metrics.

## Supporting information

NativeReadyDB v3 (634 proteins, CC-BY 4.0)

## Data Availability

NativeReadyDB v3 (634 proteins, JSON) and per-protein evidence text are released under CC-BY 4.0 in the GitHub repository at https://github.com/brhanufen/nativeready. The embedding cache (esm2_embeddings_634.npy, 3 MB) and per-paper extraction provenance (PMCID, evidence paragraph, schema-mapped record) are included. A persistent Zenodo DOI will be registered in a future release of this preprint to provide a citable archival snapshot.

## Code Availability

All code (extraction scripts, training pipeline, embedding pipeline, web frontend) is available at https://github.com/brhanufen/nativeready under the MIT license. The trained production model (model_v3.joblib) is included in the repository.

## Author Contributions

B.F.Z. designed the study, curated the dataset, implemented the model, deployed the web tool, and wrote the manuscript. Z.A. guided the formulation of the research questions and provided methodology guidance. Both authors reviewed and approved the final manuscript.

## Acknowledgments

We acknowledge the EuropePMC, RCSB PDB, and UniProt teams for providing the open APIs that made the data extraction possible. We thank the contributors to the Consortium for Top-Down Proteomics for the reference benchmark dataset (PXD047341) that anchors our positive set, and early NativeReady users for feedback. No external funding supported this work.

